# From coarse-grained molecular dynamics simulations to continuum models of biomolecular condensation

**DOI:** 10.64898/2026.10.14.759104

**Authors:** Anže Hubman, Tilen Potisk, Franci Merzel

**Affiliations:** Theory department, National Institute of Chemistry, Hajdrihova 19, 1001 Ljubljana, Slovenia; Faculty of Mathematics and Physics, University of Ljubljana, Jadranska 19, 1000 Ljubljana, Slovenia

**Author notes:** Electronic mail.

## Abstract

Biomolecular condensation contributes to cellular organization and the regulation of numerous biochemical processes. Capturing its dynamics on large spatio-temporal scales requires a macroscopic model that accurately describes both thermodynamic forces and molecular transport. Here, we investigate the applicability of the Cahn-Hilliard equation for modeling phase-separation dynamics in intrinsically disordered proteins. Using the low-complexity domain of FUS as a model system for biomolecular condensation, we perform coarse-grained molecular dynamics simulations to infer the free-energy functional and mobility coefficient entering the Cahn-Hilliard equation. The resulting continuum model accurately describes the growth of initial concentration fluctuations in the spinodal regime and the subsequent morphological evolution of the concentration fields during coarsening. We find that the free-energy functional is transferable across different stages of phase separation, whereas the mobility is not. During coarsening, we identify two kinetic regimes characterized by distinct mobilities, which persist even when a concentration-dependent mobility is introduced.

## I. INTRODUCTION

In living cells, liquid-liquid phase separation (LLPS) is a primary mechanism for the formation of biomolecular condensates^1^. Through LLPS, proteins and nucleic acids assemble into liquid-like, membraneless compartments^2^. Nearly two decades^3^ of intense research have established the important roles these condensates play in numerous cellular processes. For example, they contribute to spatiotemporal control within cells^4^, reduce stochastic noise in protein concentration levels^5^, affect RNA stability^6^, and respond to stress signals^7^, thereby contributing directly to cell fate decisions.

Intrinsically disordered proteins (IDPs) and proteins containing disordered regions (IDRs), such as FUS^8^ and Tau^9^, are particularly prone to LLPS. Their sequences can typically be described as comprising *sticker* regions that engage in transient, multivalent interactions and *spacer* regions that modulate chain flexibility^10^. Although these sticker-mediated interactions are individually weak, they collectively determine the thermodynamics and kinetics of condensate formation.

A comprehensive understanding of LLPS requires connecting molecular-scale interactions to emergent macroscopic phase behavior, making it inherently a multiscale problem^11,12^. Experimental techniques, including NMR^13^, SAXS^14^, and FRET^11^, provide important insights into the molecular structure and dynamics of proteins in both dense and dilute phases of condensates. To complement and interpret these experiments, molecular dynamics (MD) simulations play a crucial role in deciphering molecular interactions driving biomolecular condensation^15^. Atomistic MD simulations offer the most detailed molecular resolution but remain computationally demanding. Their application is restricted to relatively small systems, pre-equilibrated dense phases, and limited simulation timescales^11,16,17^.

Coarse-grained (CG) models address these limitations by reducing molecular resolution and approximating effective interactions that govern phase behavior. This enables simulations of larger systems and longer timescales than are accessible to atomistic simulations. A range of CG models has been developed for simulating biomolecular phase separation and conformational ensembles of IDPs. Examples include the explicit-solvent model Martini^18,19^ and implicit-solvent models such as HPS^20^, Mpipi^21^, and CALVADOS^22^.

Using these coarse-grained models, substantial effort has been devoted to characterizing the conformational properties of individual disordered protein chains and the equilibrium properties of biomolecular condensates^20,23^. Coexistence simulations in slab geometries are now routinely used to construct phase diagrams for different sequences^21^, characterize the interfacial properties of condensates^24^, and investigate molecular exchange between the coexisting phases^25^. Furthermore, extensive simulations across sequence space, combined with active learning, have enabled the training of machine-learning models that predict the phase behavior of IDPs directly from their sequences^26^. Despite these advances in understanding the thermodynamics of LLPS using MD simulations, the kinetics of biomolecular condensation has received considerably less attention.

Important exceptions, focusing on phase separation in the nucleation regime, include the works of Li et al.^27^ and of Sepehri and Zerze^28^. In both studies, the authors determined the nucleation free-energy barriers. While the height and shape of the nucleation free-energy barrier do not fully specify the kinetics of LLPS, they provide valuable insight into the underlying mechanism and allow for comparisons among different sequences. In a recent study, Zhang and Chu^29^ also used CG-MD to examine the roles of intermolecular contacts and salt concentration in the kinetics of phase separation.

In addition to MD simulations, continuum models offer a natural and computationally tractable approach for studying the kinetics of LLPS and subsequent morphological evolution. One such model, commonly adopted in theoretical studies^30–32^, is the Cahn-Hilliard (CH) equation, which describes phase separation through conserved diffusive dynamics of a chosen order parameter, typically the concentration field^33,34^. In its deterministic form, the CH equation describes the growth of fluctuations in the spinodal regime, where no free-energy barrier to phase separation exists. By including stochastic thermal noise, this description (the Cahn-Hilliard-Cook model^35^) can be extended to the nucleation regime, where fluctuations are required to drive the system across the nucleation barrier. Both versions of the CH model can also be used to describe the late stage droplet coarsening regime.

Despite their utility, continuum models face two key challenges. First, their validity is often assumed rather than established, which calls for systematic tests of the range of scales and conditions over which they remain valid. Second, continuum models typically contain several phenomenological parameters. Within the CH framework, these include derivatives of the free-energy functional and the mobility coefficient. Molecular dynamics simulations provide a route to both testing the underlying assumptions of these models and determining phenomenological parameters directly from the molecular system under consideration. An important step in this direction, with direct relevance to our study, was made by Silalahi et al.^36^, who developed a dynamical density functional theory (DDFT) describing the condensation of Ddx4 protein. The DDFT framework, parameterized using experimental data, was then applied to study the coalescence of protein droplets. However, their study did not assess whether the model also accurately describes earlier stages of LLPS.

In this work, we systematically investigate the applicability of the Cahn-Hilliard equation to the kinetics of biomolecular phase separation. We use coarse-grained molecular dynamics (CG-MD) simulations of the low-complexity domain of FUS (FUS-LCD) to determine both the free-energy functional and the mobility coefficient entering the Cahn-Hilliard equation. We then assess the ability of the resulting continuum model to reproduce the phase-separation kinetics observed in the MD simulations. We find that the model accurately describes initial concentration fluctuations in the spinodal regime, as well as intermediate-stage domain coarsening, with the phenomenological parameters entering the Cahn-Hilliard equation depending on the kinetic regime being described.

## II. METHODS

### A. Cahn-Hilliard equation

The Cahn-Hilliard^33,34^ equation provides a continuum description of phase separation through the temporal evolution of a conserved order parameter, here represented by the concentration field *ϕ*. Throughout this work, concentrations are expressed as volume fractions. The field dynamics are governed by gradients of the chemical potential derived from the free-energy functional and are given by:

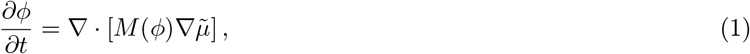

where *M* (*ϕ*) is the concentration-dependent mobility coefficient and:

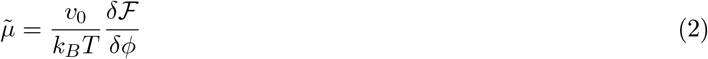

is the chemical potential, defined as the functional derivative of the free-energy functional ℱ [*ϕ*], *v*_0_ is the volume of diffusing object (average protein residue or lattice site volume) and *T* is the temperature. The mobility determines the timescale of composition evolution, while the free-energy functional defines the thermodynamic driving force for phase separation. By introducing the flux 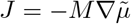, Eq. (1) takes the form of a continuity equation:

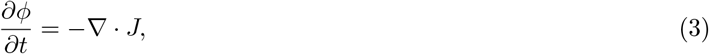

which ensures conservation of the total concentration in a closed system. A common ansatz^37^ for *M* (*ϕ*), which is also used in this work, is:

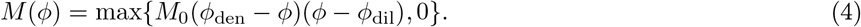

This form ensures that the flux vanishes in the coexisting dense and dilute phases, with concentrations *ϕ*_den_ and *ϕ*_dil_, respectively. We note that the DDFT framework in Ref.^36^ can be recast as a CH-type equation with the mobility *M* (*ϕ*) = *M*_0_*ϕ* by replacing particle densities *ρ* with volume fractions *ϕ* and setting *M*_0_ = *DN*, where *D* is the diffusion coefficient and *N* is the number of diffusing molecular objects (protein residues).

### B. Free-energy functional

The free-energy functional entering the Cahn-Hilliard equation is written in the square-gradient form^33^:

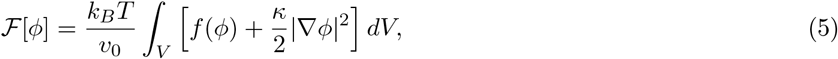

where *f* (*ϕ*) is a dimensionless bulk free-energy density and *κ* is the square-gradient coefficient. The gradient contribution accounts for the energetic cost associated with spatial variations in *ϕ*, such that a large value of *κ* corresponds to a broader interface. The integral in Eq. 5 is taken over the system volume *V*.

In this work, the bulk free-energy density is chosen to have the Flory-Huggins form^38,39^:

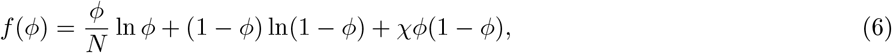

where *χ* is the Flory-Huggins interaction parameter. Equation (6) assumes that the solvent occupies a single lattice site. Although the Flory-Huggins free-energy density is a mean-field approximation, we assume that it provides a reasonable description for the nearly uncharged FUS-LCD sequence, for which long-range electrostatic interactions are expected to play only a minor role.

### C. Parameterization of the Flory-Huggins model

To parameterize the Flory-Huggins free-energy density for a given sequence and the underlying force field used in the MD simulations, the parameters *v*_0_ and *χ* must be determined. To define *v*_0_, we assume that each protein residue is represented by a hard sphere occupying a single lattice site. Each residue *k* has a diameter *σ*_*k*_, corresponding to the residue-specific Wang-Frenkel interaction parameters^40^. The associated monomer volume is therefore:

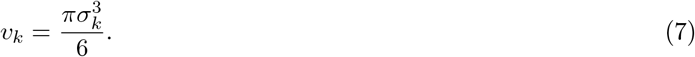

The distribution of *v*_*k*_ for the FUS-LCD sequence is given in the Supplementary Information. We define *v*_0_ as the average monomer volume:

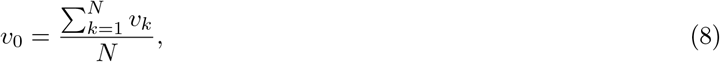

where the sum extends across all residues in the protein sequence. Therefore, the relation between the number concentration of protein residues *c* and volume fraction *ϕ* is *ϕ* = *v*_0_*c*.

The interaction parameter *χ* is determined from the coexistence concentrations *ϕ*_den_ and *ϕ*_dil_, which are obtained from slab simulations (see Sec. II G and the Supplementary Information). At equilibrium, the chemical potentials 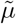 and osmotic pressures Π of the coexisting phases are equal:

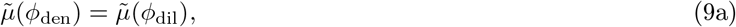

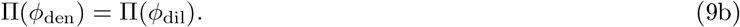

The chemical potential and osmotic pressure are defined as:

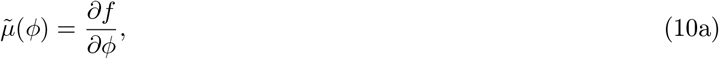

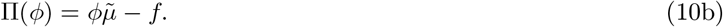

Note that Eq. 10a follows from Eq. 5 in the homogeneous limit, where ∇*ϕ* = 0, as is the case in the bulk phases away from the interface.

Using the Flory-Huggins free-energy density, these coexistence conditions yield two independent estimates of the interaction parameter:

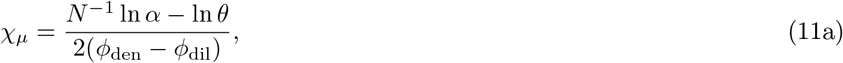

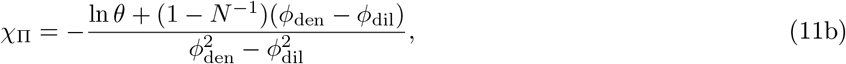

where we have introduced shorthand notation *α* = *ϕ*_den_*/ϕ*_dil_ and *θ* = (1 − *ϕ*_den_)*/*(1 − *ϕ*_dil_). We estimate *χ* using both expressions and compare the resulting values as a consistency test for the applicability of the Flory-Huggins description to the simulated system.

### D. Determination of the square-gradient coefficient

The square-gradient coefficient *κ* is determined from the relation between the surface tension *γ* and the equilibrium concentration profile:

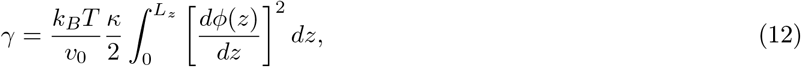

where the factor of 1*/*2 accounts for the presence of two interfaces in the slab geometry. The derivative *dϕ/dz* is evaluated numerically from the equilibrium concentration profile.

The surface tension *γ* is obtained from the slab simulations using the mechanical definition based on the pressure tensor^41^:

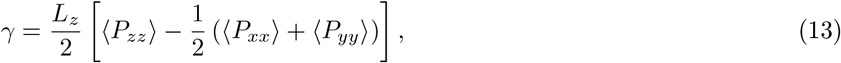

where *L*_*z*_ is the box length in the direction perpendicular to the interface and ⟨·⟩ denotes a temporal average over the simulation trajectory. The factor of 1*/*2 again accounts for the two planar interfaces in the slab geometry. The uncertainty in *κ* is estimated as standard error of the mean from four independent slab simulations.

### E. Spinodal decomposition

For a uniform mixture quenched into the spinodal regime, linearized Cahn-Hilliard theory predicts the time dependence of concentration fluctuations in reciprocal space through the temporal evolution of the structure factor *S*(**k**, *t*). At short times, this dependence is given by:

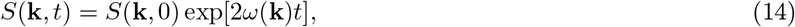

where *ω*(**k**) is the wavevector-dependent growth rate, which is positive for |**k**| *< k*_*c*_ and negative otherwise. Here, *k*_*c*_ is the critical wavevector.

To obtain the growth rate from MD simulations, we prepared elongated simulation boxes with dimensions 15 nm× 15 nm ×300 nm and *ϕ* = 0.0746. First, the system was equilibrated at *T* = 400 K for 50 ns. The temperature was then quenched to *T* = 300 K, and the system was followed for an additional 50 ns.

By design, phase separation proceeds primarily along the *z*-direction. We therefore model LLPS as a quasi-one-dimensional process, such that only the *k* = *k*_*z*_ component of **k** is considered, with an average over the *x* and *y* directions implied. Crucially, periodic boundary conditions (PBCs) restrict the set of allowed wavevectors to:

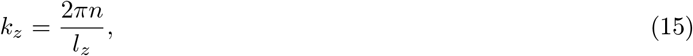

where *n* is an integer. Using a relatively long simulation box along the *z*-direction therefore allows the system to develop a significant number of unstable modes with *ω*(*k*) *>* 0. To obtain *ω*(*k*), we first computed the time-dependent structure factor from the particle positions using:

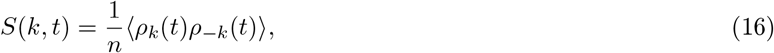

where *ρ*_*k*_(*t*) is the Fourier component of the instantaneous density field, *n* is the total number of residues, and ⟨… ⟩ denotes an ensemble average over 30 independent replicas. The growth rate *ω*(*k*) was then obtained by linear regression of log[*S*(*k, t*)*/S*(*k*, 0)] versus *t*.

### F. Phase separation dynamics in the coarsening regime

To investigate phase-separation dynamics during the coarsening stage and determine the mobility coefficient, we developed an approach inspired by the works of Silalahi et al.^36^ and Barclay & Lucas^42^. The method presented here is also similar to the nonequilibrium approach we introduced recently for computing thermal conductivities in liquids^43^.

First, an elongated simulation box with dimensions *l*_*x*_ = 30 nm ×*l*_*y*_ = 30 nm ×*l*_*z*_ = 100 nm was created. A central slab of width *h* = 40 nm was then selected, and proteins were placed randomly within this region using a combination of <monospace>polyply</monospace>^44^ and several in-house Python scripts until the concentration reached a prescribed value *ϕ*_0_ ≈0.1 (*ϕ*_0_ *< ϕ*_den_). To equilibrate the system and prevent the proteins from leaving the selected region, we introduced a flat-bottom restraint:

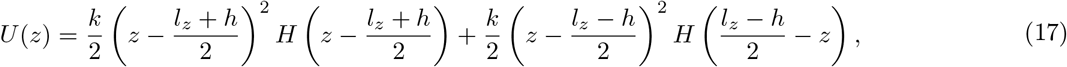

where *H*(*s*) is the Heaviside function and *k* = 1.0 kcal mol^*−*1^ Å^*−*2^ is the harmonic force constant. The system was equilibrated for 50 ns at *T* = 360 K. The restraint was then removed, and the temperature was quenched to *T* = 300 K. The system was subsequently allowed to evolve for 200 ns until equilibrium was established. This protocol was repeated for 15 independent replicas.

There are two key differences between our method and that of Barclay & Lucas. First, in Ref.^42^, the central region is prepared at the equilibrium concentration, and the width of the interface is perturbed to study interfacial relaxation. In contrast, in our approach, the central region is prepared at a concentration below the equilibrium value, such that both densification of the central region and reshaping of the interface are observed. Second, in Ref.^42^, the interface is perturbed by performing Monte Carlo swap moves that exchange particles between the dense and dilute regions, thereby imposing a concentration gradient across the interface. While elegant, this approach is feasible only when the particles are small compared with the characteristic interfacial width and is therefore not easily transferable to polymeric systems. In our approach, the initial condition is instead generated using a flat-bottom potential. Although we use a simple flat-bottom potential in the present work, this approach could in principle be adapted by tailoring the potential to the specific system and desired initial condition.

To inspect the dynamical evolution of the system, we computed the concentration profiles *ϕ*(*z, t*) as a function of time along the *z*-axis. The profiles were obtained using simple histograms with *n*_*z*_ = 256 bins. We did not smooth the histograms or use kernel density estimators in order to avoid artificially broadening the interface. Because the nonequilibrium initial condition is reproducibly defined, apart from thermal fluctuations, the concentration profiles at each time point were calculated independently for each replica and then averaged across replicas. Thus, the reported *ϕ*(*z, t*) represents the ensemble-averaged concentration profile at each time *t*.

To determine the mobility coefficient that best describes the time evolution of the concentration profiles obtained from MD simulations, we numerically propagated the Cahn-Hilliard equation from the MD-derived concentration profile at *t* = 0 for different values of the mobility. The optimal mobility was determined by minimizing the loss function ℒ, which quantifies the deviations between the predicted and MD-derived concentration profiles:

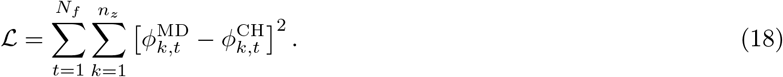

Here, the sums extend over all *N*_*f*_ considered frames and *n*_*z*_ bins. The uncertainty in the estimated mobility was determined using jackknife resampling. To obtain numerical solutions of the CH equation, we developed a custom solver implemented using a spectral method^37^.

## G. Molecular dynamics simulations

Molecular dynamics simulations were performed using LAMMPS (version 30Mar2026).^45^ We used the coarse-grained model Mpipi-GG^46^, which is a modified parameterization of the original Mpipi model^21^. In Mpipi-GG, amino-acid residues are represented by single beads connected by harmonic springs. The nonbonded interactions consist of the Wang-Frenkel^40^ potential and a screened electrostatic contribution described by the Debye-Hückel potential. All production trajectories after the quench were performed at *T* = 300 K using an inverse Debye screening length of 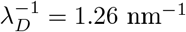 nm^*−*1^, corresponding to an ionic strength of 0.15 M for a monovalent salt. The equations of motion were integrated using a timestep of 20 fs. The solvent was treated implicitly using a Langevin thermostat as implemented in the <monospace>fix langevin</monospace> command in LAMMPS^47,48^. In all production simulations, the damping time of the thermostat was set to *τ* = 25 ps in equilibrium simulations and to *τ* = 5 ps in nonequilibrium runs.

## H. Fine-tuning the Mpipi-GG model

Preliminary slab simulations showed that, for FUS-LCD, the original Mpipi-GG force field predicts coexistence concentrations that differ from experiment. To improve the agreement, we fine-tuned the force field specifically for the FUS-LCD sequence.

Following a commonly used approach^18,49^, the depth (*ϵ*) of all protein-protein interaction potentials was uniformly scaled by a factor *α <* 1. The scaling parameter was adjusted to reproduce the experimentally determined transfer free energy between the dilute and dense phases:

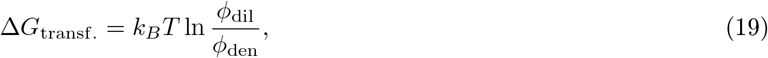

where Δ*G*_transf._ was computed from experimentally reported coexistence concentrations.

## III. RESULTS AND DISCUSSION

### A. Fine-tuned Mpipi-GG model

First, we assess the ability of the original Mpipi-GG force-field parameterization to reproduce the experimentally determined phase behavior of FUS-LCD. The model predicts a dense-phase concentration of *ϕ*_den_ ≈ 0.31 and an extremely low dilute-phase concentration of *ϕ*_dil_ ≈ 0 (results not shown). Because the dilute-phase concentration is very small, it cannot be determined reliably for the simulation size and timescale considered here.

To determine an appropriate value of the scaling parameter *α*, we fitted the dependence of Δ*G*_transf._ on *α* using a linear relationship (see Fig. 1) and a weighted least-squares method, with the inverse variances of Δ*G*_transf._ used as weights. The resulting fit was used to estimate the value of *α* corresponding to Δ*G*_transf._ ≈ −11 kJ/mol, yielding

**FIG 1.**
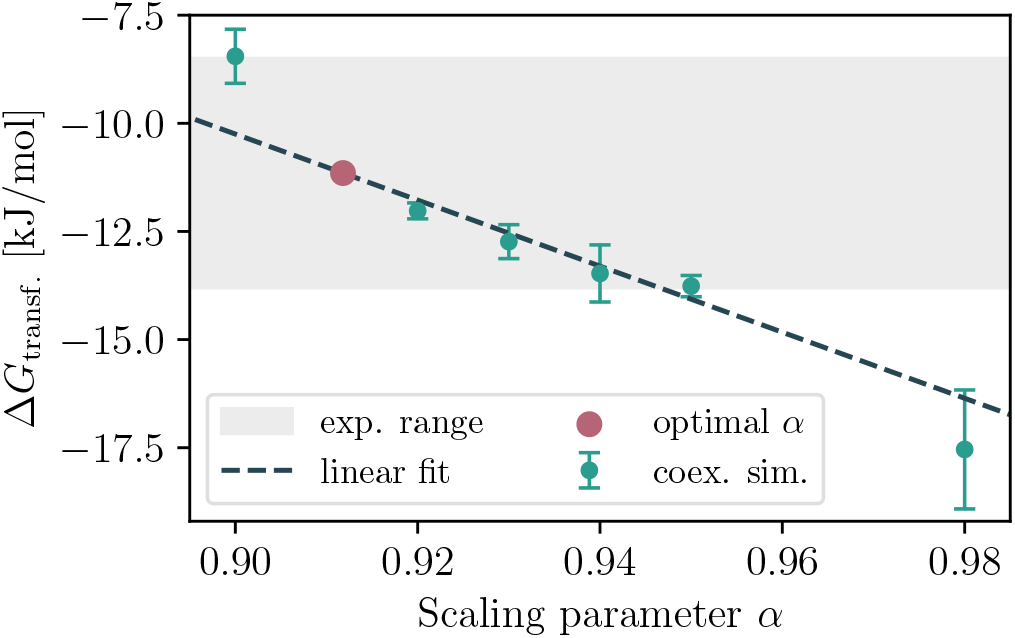
Dependence of the transfer free-energies on the scaling parameter *α*. Error bars represent the standard deviations computed across 5 blocks. Experimental range of the transfer free-energies (shaded area) is taken from Ref^18^.

*α*≈ 0.91. This value was selected because it corresponds to a transfer free energy near the center of the range first computed by Benayad et al.^18^, based on experimental measurements^50–53^. We emphasize that this empirical correction is unlikely to be transferable to other sequences. Furthermore, a uniform scaling assumes that the *relative* abundances of specific residue-residue contacts are already correctly captured by the unmodified force field.

Using the molar mass of FUS-LCD (*M*_FUS*−*LCD_ = 17.168 kg/mol^18^) the coexistence volume fractions can be converted to mass concentrations, yielding *ρ*_den_ ≈356 g/L and *ρ*_dil_ ≈3 g/L. These values are consistent with the experimentally reported coexistence concentrations at *T* = 300 K. For example, dilute-phase concentrations for FUS-LCD have been reported in the range 0.2 −4.0 g/L^51,54,55^, while dense-phase concentrations range from 320 −447 g/L^53,54^. This agreement is reassuring because matching the transfer free-energies optimizes the log-ratio of the coexistence concentrations rather than the *individual* concentrations directly. All subsequent simulations were performed using the modified force field with *α* = 0.91.

### B. Thermodynamics

Next, we discuss the parameterization of the free-energy functional. We begin with the estimation of the interaction parameter *χ* and the assessment of the validity of the Flory-Huggins model. Using coexistence concentrations measured in four independent slab simulations, we obtain *χ*_*µ*_ = 0.629±0.003 and *χ*_Π_ = 0.614 ± 0.001 from the equality of chemical potentials 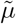 and osmotic pressures Π, respectively. These two estimates differ by approximately 2%, which indicates a small but systematic discrepancy. However, given the approximations involved in the Flory-Huggins model, we consider this discrepancy to be reasonable. To avoid ambiguously selecting between these two estimates and to minimize the error in the predicted binodal concentrations, we instead determined the optimal *χ* by minimizing the cost function ℒ:

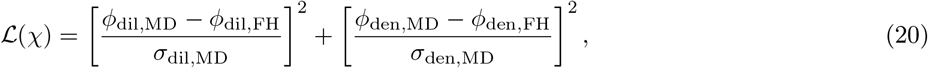

where the subscript ‘MD’ denotes concentrations obtained from the slab simulations, with the corresponding standard errors *σ*, while the subscript ‘FH’ denotes concentrations predicted using the Flory-Huggins model for a given *χ*. This approach yields *χ*_opt_ ≈0.614 (see SI for details), and we therefore use this value in the remaining analyses. Nonetheless, it is important to note that this self-consistency test only demonstrates that the Flory-Huggins model provides a good description of the binodal concentrations. It does not, however, guarantee that the curvature of *f* (*ϕ*), which is important for an accurate description of the dynamics, is correct over the entire concentration range. We return to this issue in the next two sections.

The second contribution to the free-energy functional is the squared-gradient term, 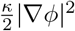. As discussed in Section II D, we determined *κ* from the surface tension *γ* and the equilibrium concentration profile *ϕ*(*z*). Using Eq. 13, we obtain a surface tension of *γ* = 98 ± 22 *µ*N/m, which is consistent with the range reported for FUS-LCD droplets described by the modified Martini2 force field^18^ and falls within the experimental range reported by Taylor et al.^56^ for the LAF1 and Whi3 sequences. Using this estimate of *γ* in Eq. 12 together with the equilibrium concentration profiles, we obtain *κ* = 84 ± 17 *Å*^2^.

### C. Spinodal decomposition

A homogeneous mixture quenched into the spinodal regime undergoes rapid spontaneous phase separation. Our parameterization of the Flory-Huggins model predicts a spinodal region spanning *ϕ* = 0.034 to *ϕ* = 0.147, placing our choice for the initial concentration *ϕ*_0_ = 0.0746 well within the spinodal regime. For spinodal decomposition initiated from a uniform initial concentration *ϕ*_0_, linearized Cahn-Hilliard theory predicts the *k*-dependent growth rate *ω*(*k*):

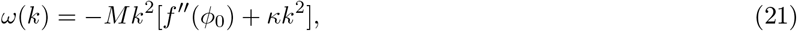

where *M* = *M* (*ϕ*_0_) is the mobility and *f*^*′′*^(*ϕ*_0_) is the second derivative of the free-energy density evaluated at *ϕ*_0_. This analytical expression provides a means to assess the validity of Cahn-Hilliard theory and to estimate the parameters *χ, M*, and *κ* by fitting the theoretical *ω*(*k*) to the values obtained from MD simulations, as described in Section II E. The time-dependent structure factors obtained from the MD simulations and used to compute *ω*(*k*) are shown in the Supplementary Information.

To estimate the unknown parameters (*θ*), given the observations *D*= [*ω*(*k*_1_), …, *ω*(*k*_*n*_)] we used Bayesian inference. Bayes’ theorem states that:

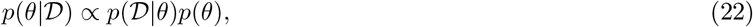

where *p*(*θ*|D) is the posterior distribution, *p*(D|*θ*) is the likelihood and *p*(*θ*) is the prior. We used flat priors (*p*(*θ*) = const.) and a Gaussian likelihood corresponding to a chi-squared objective function. The correlations among different *k*-points were neglected, such that the covariance matrix was assumed to be diagonal. We performed two inference procedures, keeping *χ* fixed at the value determined in Section III B in both cases. In option 1, *κ* was fixed at the mean value obtained from equilibrium slab simulations, and only the mobility coefficient *M* was inferred (*θ* = *M*). In option 2, posterior distributions were obtained for both *M* and *κ*, so that *θ* = (*M, κ*). The results are shown in Fig. 2.

**FIG 2.**
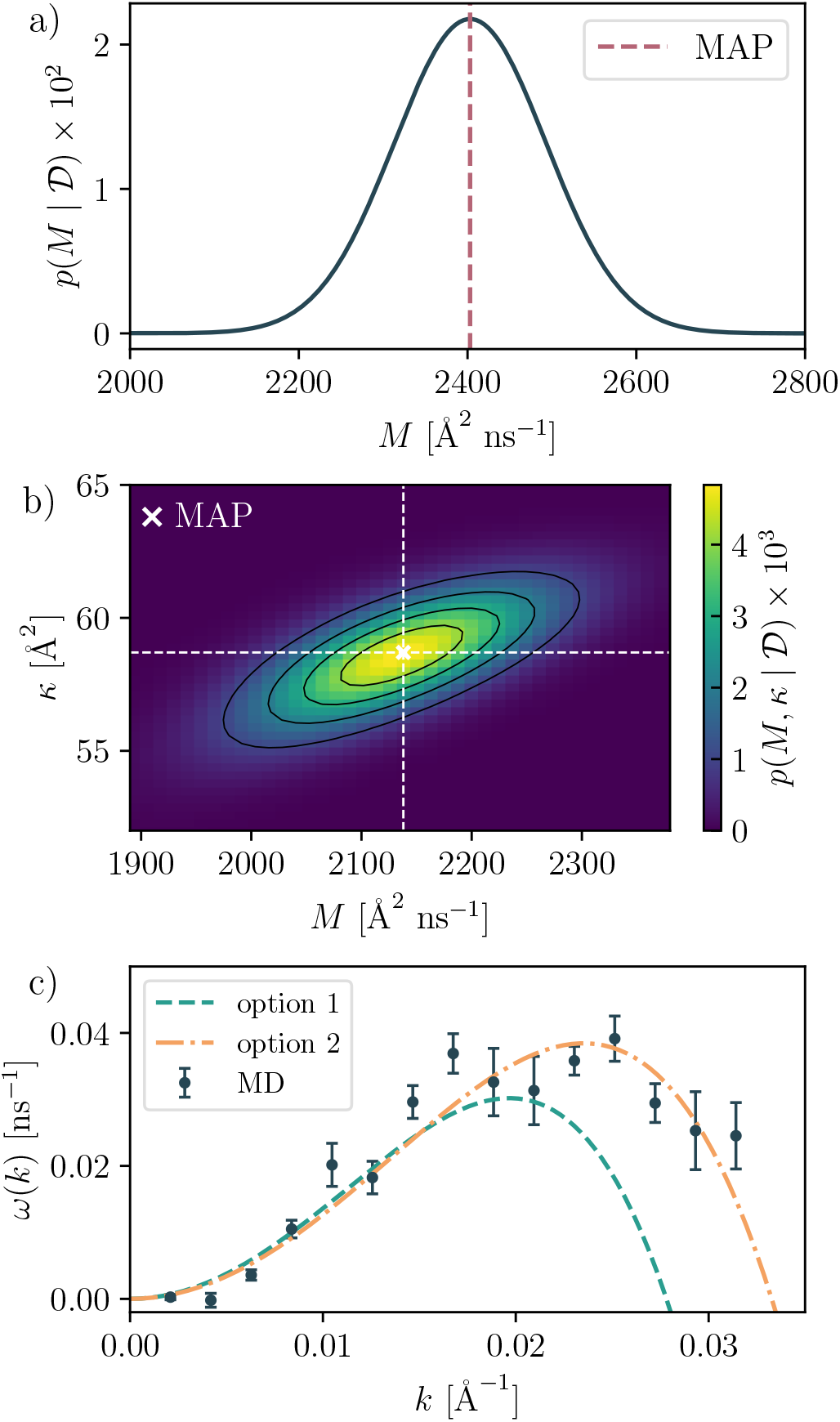
(a) Posterior distribution of the mobility coefficient estimated in option 1. (b) Joint posterior distribution of the mobility coefficient and squared-gradient coefficient estimated in option 2. (c) Growth rates obtained from MD simulations (black dots) and model predictions based on the estimates from options 1 (green line) and 2 (orange line).

As shown in Fig. 2a and Fig. 2b, we obtain relatively consistent estimates of the mobility coefficient, with the maximum-a-posteriori (MAP) estimates differing by roughly 15%. A larger discrepancy is observed between the two estimates of *κ*, which differ by approximately 30%. This discrepancy can be rationalized by two factors. First, in option 1, the estimate of *κ* is based on the surface tension, which is a noisy quantity that is difficult to converge in MD simulations (see the SI for details). Second, jointly inferring *M* and *κ* from the MD-derived *ω*(*k*) in option 2 partly inherits inaccuracies in the Flory-Huggins free-energy density, which are subsequently propagated into the parameter estimates.

With these discrepancies in mind, the results shown in Fig. 2c support the validity of Cahn-Hilliard theory for describing the growth rate during the early stages of spinodal decomposition. This is also consistent with the recent work of Zhang et al.^57^, who arrived at a similar conclusion using a different metric to describe concentration fluctuations. In our case, the agreement is particularly good for the lowest 10 wavevectors. For the remaining portion of the *k*-axis, the agreement is qualitative. However, it should be kept in mind that the exact position of the maximum growth rate and the critical *k*-vector are extremely sensitive to the values of *χ* and *κ*.

We conclude this section with a remark on the quasi-1D treatment of phase separation. As can be seen in Fig. 2c, only the lowest 15 wavevectors were used to compute *ω*(*k*). This quasi-1D treatment along the *z*-direction is justified only for wavevectors *k*_*i*_ whose characteristic length *l*_*i*_ = 2*π/k*_*i*_ is larger than the simulation box dimensions in the *x*- and *y*-directions. In our case, the largest *k* corresponds to *l*≈ 1.33*l*_*x*_ = 1.33*l*_*y*_, thus approaching the limit of applicability. If this condition is not satisfied, the dynamics in directions orthogonal to the *z*-direction cannot be neglected. This implies that the MD-derived *ω*(*k*) is more reliable at smaller *k*. The same condition therefore limits the largest *k* at which *ω*(*k*) can be estimated, as accessing larger *k* would require reducing the box dimensions in the *x*- and *y*-directions. However, these dimensions cannot be reduced arbitrarily without introducing substantial artifacts in chain behavior due to the periodic boundary conditions.

### D. Coarsening dynamics

Next, we investigated whether the Cahn-Hilliard equation can also describe the evolution of concentration profiles during the coarsening stage, that is, when the system leaves the early spinodal decomposition regime and interfaces begin to form. In one dimension, this typically corresponds to the formation of several localized domains, which later diffuse together and merge. We show an example of such a pattern in the Supplementary Information. Our MD setup can therefore be understood as an idealization in which one of these domains is isolated so that its time evolution can be studied independently. We tested two models: in one case, we used constant mobility, while in the other, the mobility was concentration dependent, as given by Eq. 4.

In Fig. 3, we show examples of concentration profiles at different times computed from MD simulations. Along with the mean 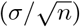 profile, computed over *n* replicas, we also plot the standard deviation (*σ*) and the standard error of the mean (*σ/ n*). The former characterizes the magnitude of thermal fluctuations and converges to a finite value as *n* → ∞.

**FIG 3.**
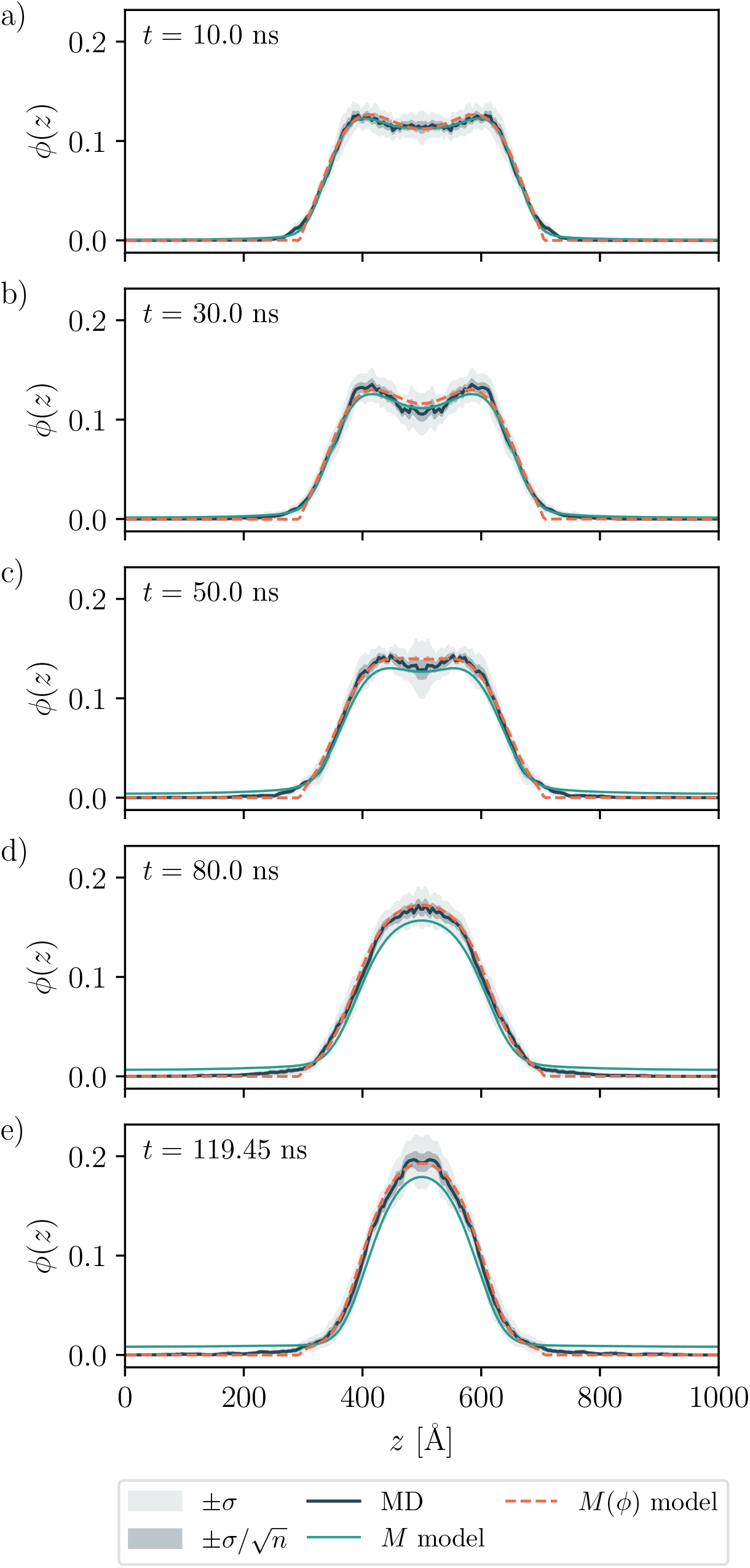
(a)-(e) Concentration profiles at different stages of phase separation obtained from MD simulations (black), together with predictions using constant mobility (green) and concentration-dependent mobility (orange dashed line). The light gray region indicates the standard deviation, reflecting thermal fluctuations and, strictly speaking, shot noise from histogramming, while the dark gray region indicates the standard error of the mean.

The latter instead converges to zero → ∞ as *n* and can therefore be used to assess systematic discrepancies between the MD simulations and the continuum model. Qualitatively, the MD-derived concentration profiles show a characteristic evolution in which the central region first develops a dip, followed by interfacial relaxation and densification of the central region until equilibrium is established. This evolution is broadly consistent with that observed by Silalahi et al.^36^, who predicted similar behavior using their DDFT model for a different protein sequence.

Let us first consider the Cahn-Hilliard model with constant mobility. The predictions are shown in green in Figs. 3a-e. As can be seen, very good agreement with the MD simulations is obtained using this simple model. This suggests that both the form of the Cahn-Hilliard equation and the parameterization of the free-energy functional already capture the essential field dynamics, with the mobility coefficient primarily determining the absolute timescale.

One limitation of the constant-mobility model is that it overestimates the rate at which mass is transported out of the initially dense region. This becomes apparent at longer times (see Figs. 3d and 3e). In addition, the model predicts a slightly different equilibrium dilute-phase concentration than the MD simulations. Because the total mass is conserved, this difference necessarily leads to a corresponding difference in the equilibrium dense-phase concentration. The resulting slight underestimation of the dense-phase concentration therefore does not indicate a failure of the model itself. In principle, a rough correction can be made in which the excess mass in the dilute phase is redistributed equally among bins in the dense region of the concentration profile. These discrepancies are largely resolved by introducing a concentration-dependent mobility (see orange dashed lines in Figs. 3a-e). The rate of mass transport into the dilute phase is reduced, and the dense-phase concentration at equilibrium (*t* ≈ 120 ns) agrees closely with the MD simulations.

In the preliminary analyses, we observed that our Cahn-Hilliard model correctly reproduces the *sequence* of morphologies observed in the MD simulations. However, we found that a single value of the mobility *M* (or the prefactor *M*_0_ in the concentration-dependent case) provides poor fits over the entire coarsening regime. We therefore divided the MD data into 24 blocks of equal length and determined the optimal mobility for each block separately. This analysis revealed that the domain relaxation proceeds in roughly two kinetic regimes, connected by a smooth transition (see Fig. 4). These two regimes are described by two distinct mobilities. Crucially, using a concentration-dependent mobility does not eliminate these two regimes. We find that the transition between the two regimes coincides with the point at which the initial concentration dip relaxes to the vicinity of its initial value, after which the densification of the domain proceeds.

**FIG 4.**
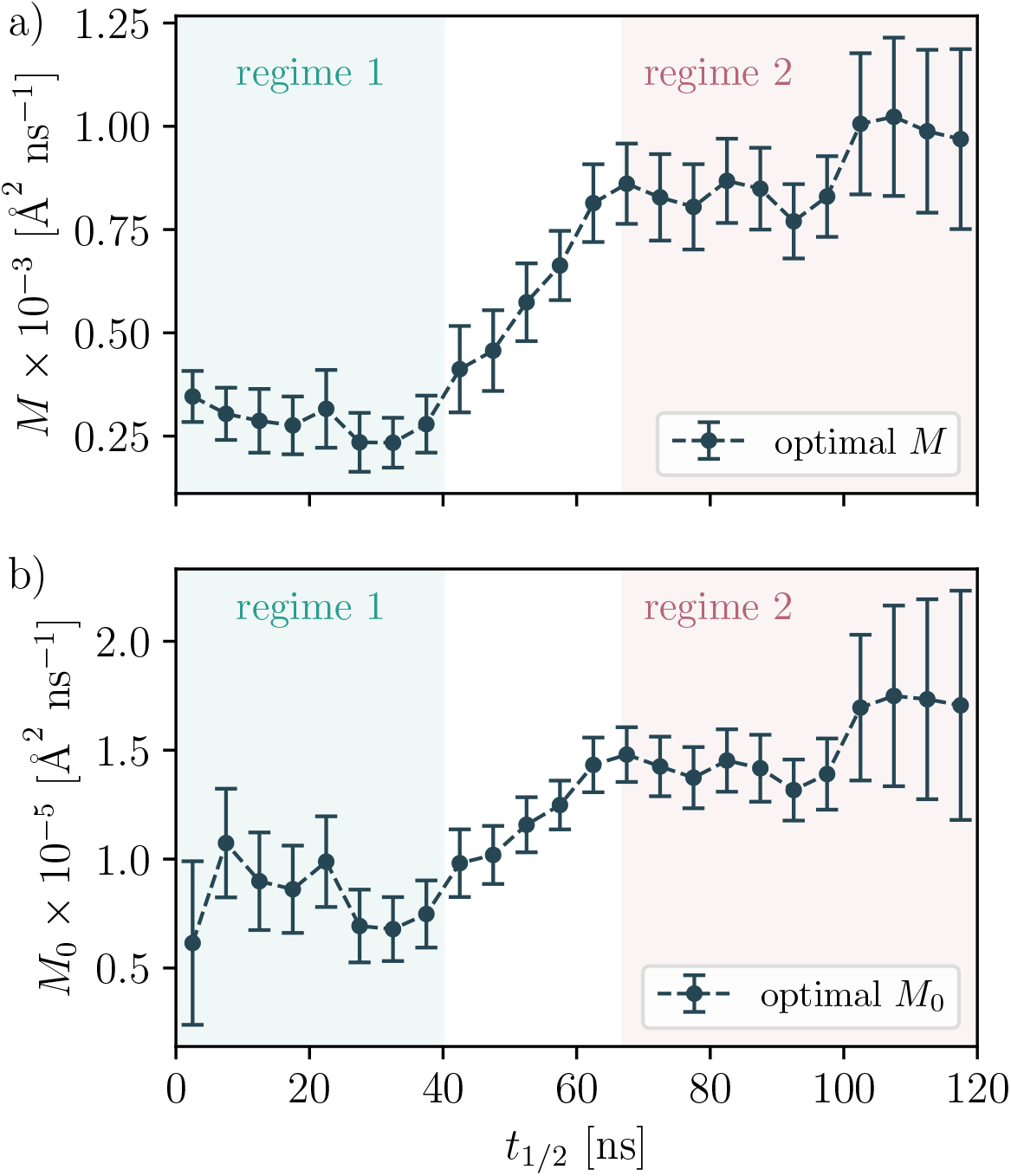
(a) Optimal constant mobility coefficient at different stages of phase separation. (b) Optimal mobility prefactor for the concentration-dependent mobility model at different stages of phase separation. The concentration-dependent mobility spans the range 0–2000 *Å*2ns^*−*1^, which brackets the range obtained for the constant-mobility model. The upper bound is also close to the mobility obtained in the spinodal regime.

The emergence of different kinetic regimes can be rationalized in two ways. First, as the initial concentration profile with steep interfaces relaxes toward equilibrium, mass is transported through both bulk and interfacial diffusion (see also the discussion in Ref.^37^ and references therein). The relative contributions of these two transport mechanisms change during the relaxation, which could result in an apparent stage dependence of the effective mobility. Second, the dynamics may involve at least one additional slow degree of freedom evolving on a timescale comparable to that of the concentration field. This would not be particularly surprising, since, for example, reconfiguration dynamics of IDPs in the dense phase of a condensate can be very slow. If such a variable is not explicitly included in the continuum description, its dynamics could manifest as a stage-dependent effective mobility.

Akin to the common approach of transforming a non-Markovian system into a Markovian one by expanding the dimensionality of the state space^58^, one could construct a multidimensional continuum model in which the concentration field is coupled to additional dynamical variables. For example, one could introduce a variable describing the relative contributions of different modes of mass transport. Identifying the relevant additional degrees of freedom and constructing such an extended model are beyond the scope of the present work and are left for future study.

## IV.CONCLUSIONS

In this work, we have demonstrated how the Cahn-Hilliard equation, applied to the phase separation of intrinsically disordered proteins, can be both validated and parameterized using MD simulations. This points to the possibility of using relatively inexpensive, but carefully designed, MD simulations to parameterize continuum models that can subsequently be used to study macroscopic behavior that would be computationally too expensive to access using MD alone. While recent advances in computational resources have made it possible to study some macroscopic phenomena using very large-scale MD simulations, such as in the work of Jung et al.^59^, these simulations remain computationally demanding. Continuum models therefore provide a complementary route to accessing macroscopic behavior at substantially lower computational cost.

A major challenge we encountered in this work is the construction of an accurate free-energy functional. This could be addressed by sacrificing some of the physical interpretability of the functional and instead making use of recent advances in machine learning. Such methods allow MD simulations to be used to train free-energy functionals and may provide a more automated procedure for learning functionals for different protein sequences. This is also an active area of research in classical density functional theory (cDFT) for complex liquids, as demonstrated, for example, by B. Cheng in a recent work^60^.

## ACKNOWLEDGMENTS

The authors acknowledge the financial support under Grant Nos. P1-0010 and J1-50033 from the Slovenian Research and Innovation Agency. We acknowledge the use of ChatGPT and Gemini in improving the quality of the text and in generating the graphical abstract accompanying this manuscript.

## AUTHOR DECLARATIONS

## Author Contributions

**Anže Hubman**: Conceptualization (equal); Formal analysis (lead); Methodology (equal); Validation (lead); Visualization (lead); Writing - original draft (lead). **Tilen Potisk**: Conceptualization (equal); Formal analysis (equal); Methodology (equal); Supervision (equal); Validation (equal); Writing - original draft (supporting); Writing - review and editing (equal) **Franci Merzel**: Conceptualization (equal); Formal analysis (equal); Funding acquisition (lead); Methodology (equal); Supervision (lead); Validation (equal); Writing - original draft (supporting); Writing - review and editing (equal).

## DATA AVAILABILITY STATEMENT

The data are available from the authors upon request.

