## Supplementary Information for "From coarse-grained molecular dynamics simulations to continuum models of biomolecular condensation"

(Dated: 14 October 2026)

### I. FASTA SEQUENCE OF FUS-LCD

```
  1 MASNDYTQQA TQSYGAYPTQ PGQGYSQQSS QPYGQQSYSG YSQSTDTSGY
 51 GQSSYSSYGQ SQNTGYGTQS TPQGYGSTGG YGSSQSSQSS YGQQSSYPGY
101 GQQPAPSSTS GSYGSSSQSS SYGGQPQSGSY SQQPSYGGQQ QSYGQQQSYN
151 PPQGYGQQNQ YNS
```

### II. ESTIMATION OF THE MONOMER VOLUME

The definition of the "monomer volume" $v_0$ is a modeling choice required to map the MD simulations onto the Flory-Huggins model[1,2]. As explained in the main text, we define $v_0$ using a sequence-averaged hard-sphere diameter, with the residue-specific Wang-Frenkel parameter $\sigma$ taken as the sphere diameter[3]. In Fig. S1, we show the distribution of the corresponding monomer volumes for the FUS-LCD sequence. As can be seen, the distribution is relatively broad. Nevertheless, the mean (0.113 $nm^3$) and median (0.106 $nm^3$) values are close to each other.

It is important to note that the parameterization of the Mpipi-GG force field[4] was performed in a temperature range around ambient temperature. Consequently, the $\sigma$ parameters should not be interpreted as defining a strict hard-sphere repulsion. Rather, they are effective parameters that, together with the other force-field parameters, capture interactions relevant to the phase behavior over the parameterization temperature range. Thus, interpreting $\sigma$ as a literal hard-sphere diameter is an approximation introduced by the mapping onto the Flory-Huggins model.

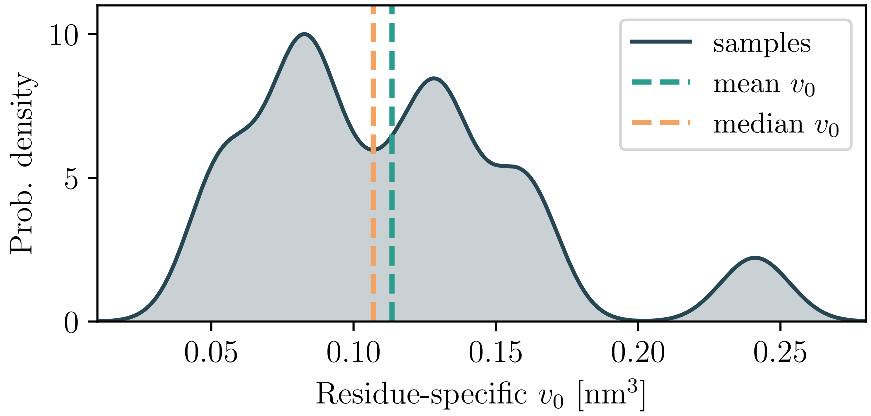


FIG. S1. Distribution of residue-specific monomer volume $v_0$ along the FUS-LCD sequence and the corresponding average (dashed line). The distribution is obtained using kernel-density estimation using a Gaussian kernel.

---

[a)] Also at Faculty of Mathematics and Physics, University of Ljubljana, Jadranska 19, 1000 Ljubljana, Slovenia.

### III. DETERMINATION OF $\phi_{\rm dil}$ AND $\phi_{\rm den}$

To determine the coexisting phase compositions, $\phi_{\rm dil}$ and $\phi_{\rm den}$, expressed as volume fractions, we analyzed the equilibrium concentration profiles obtained from long ($> 1.4$ $\mu$s) slab simulations (see Fig. S2). The profiles were averaged over all frames in each trajectory and, exploiting the symmetry of the system, also with respect to the center of the simulation box. Based on visual inspection, each profile was then divided into three regions corresponding to the dense phase, dilute phase, and interface. Estimates of $\phi_{\rm dil}$ and $\phi_{\rm den}$ were obtained by averaging over the respective regions and across all four replicas. Using the paired estimates of $\phi_{\rm dil}$ and $\phi_{\rm den}$ from each replica, we also estimated the Flory-Huggins interaction parameters using the chemical potential route ($\chi_\mu$) and the osmotic pressure route ($\chi_\Pi$); see Main Text for explicit expressions. The resulting values are reported in Table S1.

| replica | $\phi_{\rm den}$ | $\phi_{\rm dil}$ | $\chi_\mu$ | $\chi_\Pi$ |
|---|---|---|---|---|
| 1 | 0.2170 | 0.0043 | 0.6214 | 0.6143 |
| 2 | 0.2152 | 0.0023 | 0.6288 | 0.6139 |
| 3 | 0.2154 | 0.0018 | 0.6319 | 0.6140 |
| 4 | 0.2140 | 0.0033 | 0.6241 | 0.6134 |

TABLE S1. Coexistence volume fractions, $\phi_{\rm den}$ and $\phi_{\rm dil}$, and the corresponding Flory–Huggins interaction parameters obtained from the chemical potential ($\chi_\mu$) and osmotic pressure ($\chi_\Pi$) routes for four independent MD simulations.

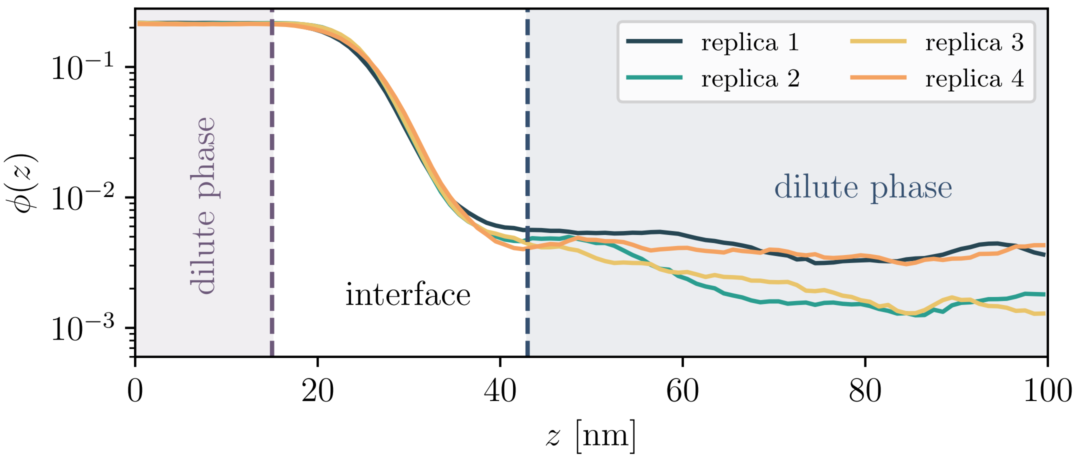


FIG. S2. Equilibrium concentration profiles from four independent MD simulations, with the corresponding regions indicated.

### IV. FINE-TUNING OF THE MPIPI-GG FORCE FIELD

To determine the optimal scaling parameter $\alpha$[5,6], we tested different values in the range $0.85 \leq \alpha \leq 1.0$, with $\alpha = 1.0$ corresponding to the unmodified force field (see Table S2). For $\alpha = 0.85$, no stable condensates were observed. For $\alpha = 1.0$, all proteins remained in the dense phase. Visual inspection of the trajectories revealed that some proteins transiently dissociated from the dense phase but were quickly reabsorbed. Therefore, the dilute-phase concentration could not be measured reliably. This behavior is consistent with an extremely low equilibrium dilute-phase concentration, for which an accurate measurement would require a substantially larger simulation box.

In this set of simulations, 100 proteins were simulated in an elongated simulation box with dimensions $l_x = 20$ nm, $l_y = 20$ nm, and $l_z = 90$ nm. Each simulation was run for at least 500 ns at $T = 300$ K.

| $\alpha$ | $\Delta G$ [kJ/mol] | $\sigma(\Delta G)$ [kJ/mol] | $\rho_{\mathrm{den}}$ [g/L] | $\sigma(\rho_{\mathrm{den}})$ [g/L] | $\rho_{\mathrm{dil}}$ [g/L] | $\sigma(\rho_{\mathrm{dil}})$ [g/L] |
|---|---|---|---|---|---|---|
| 0.90 | -8.4504 | 0.5709 | 306.4571 | 6.9082 | 11.5492 | 2.6922 |
| 0.92 | -12.0243 | 0.1684 | 361.6906 | 1.7862 | 2.9448 | 0.2006 |
| 0.93 | -12.7355 | 0.3586 | 383.5932 | 1.2153 | 2.4368 | 0.3352 |
| 0.94 | -13.4710 | 0.6024 | 400.7973 | 1.6086 | 2.0969 | 0.5566 |
| 0.95 | -13.7638 | 0.2246 | 419.3837 | 1.1197 | 1.7140 | 0.1484 |
| 0.98 | -17.5400 | 1.1229 | 465.4180 | 0.3949 | 0.6160 | 0.2264 |

TABLE S3. Transfer free energies and phase concentrations as a function of the scaling parameter $\alpha$, including the corresponding standard deviations.

| Scaling factor $\alpha$ | State | Comment |
|---|---|---|
| 0.85 | dissolved | / |
| 0.90 | stable | / |
| 0.92 | stable | / |
| 0.93 | stable | / |
| 0.94 | stable | / |
| 0.95 | stable | / |
| 0.98 | stable | / |
| 1.00 | stable | no proteins in dilute phase |

TABLE S2. Equilibrium state as a function of the scaling factor $\alpha$.

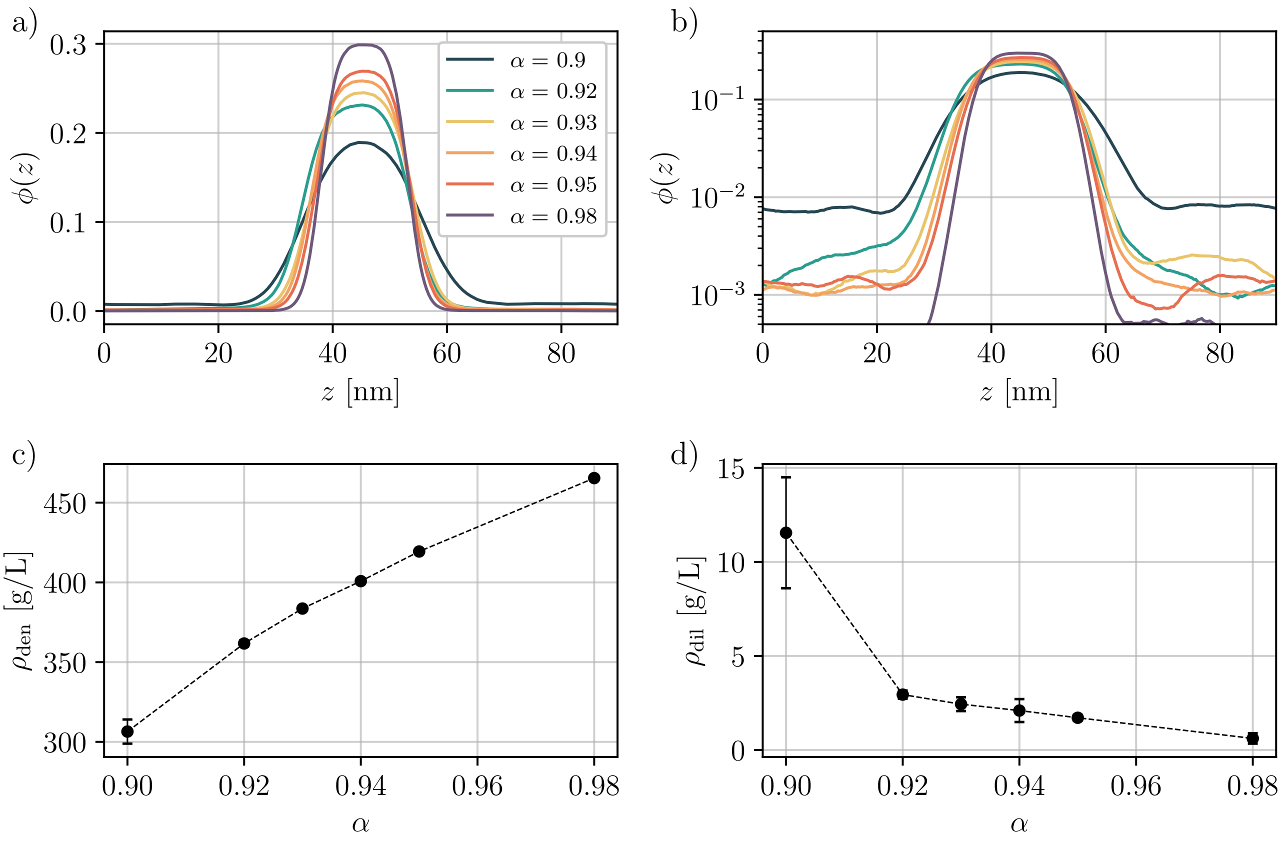


FIG. S3. (a) Time-averaged volume fraction profiles obtained from slab-coexistence simulations for different values of the scaling parameter $\alpha$. (b) The same profiles shown on a logarithmic scale to better visualize the dilute-phase volume fractions. Densities of (c) dense and (d) dilute phase, respectively, as a function of $\alpha$.

### V. ESTIMATION OF THE INTERACTION PARAMETER $\chi$

As stated in the main text and shown here in Fig. S4a, the interaction parameters estimated from the equality of chemical potentials and the equality of osmotic pressures show minor systematic deviations, which are a manifestation of the inaccuracies associated with the use of the Flory-Huggins model. To avoid ambiguously selecting between these two estimates, the optimal $\chi$ was found by minimizing the cost function $\mathcal{L}$:

$$\mathcal{L}(\chi) = \left[\frac{\phi_{\text{dil,MD}} - \phi_{\text{dil,FH}}}{\sigma_{\text{dil,MD}}}\right]^2 + \left[\frac{\phi_{\text{den,MD}} - \phi_{\text{den,FH}}}{\sigma_{\text{den,MD}}}\right]^2, \tag{S1}$$

where the subscript 'MD' denotes concentrations obtained from the slab simulations with the corresponding standard errors $\sigma$, while the subscript 'FH' denotes concentrations predicted using the Flory-Huggins model for a given $\chi$. As shown in Fig. S4b, the minimum of $\mathcal{L}$ is found at $\chi = 0.614$.

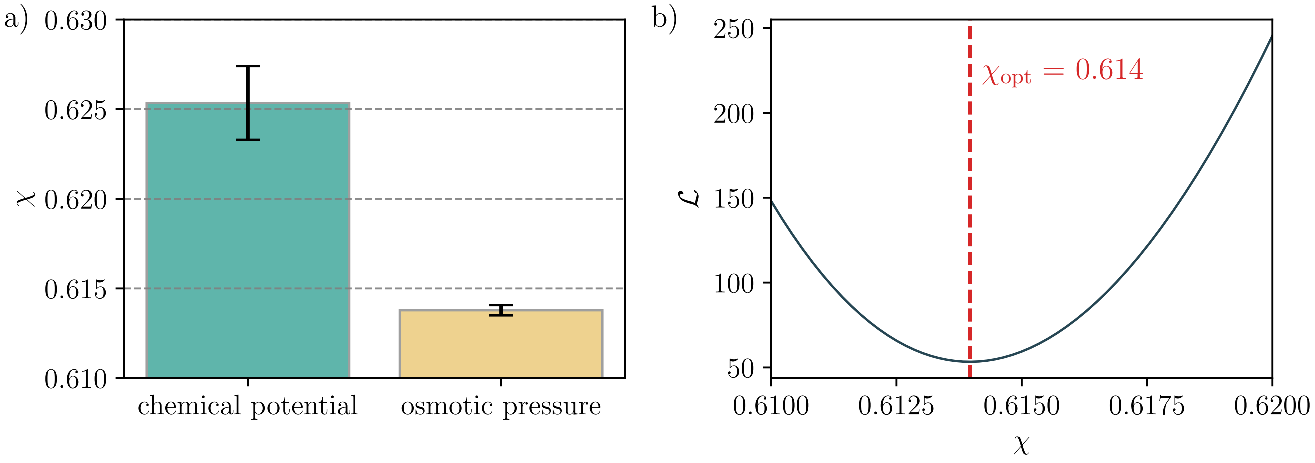


FIG. S4. (a) Estimates of $\chi$ from the equality of chemical potentials and the equality of osmotic pressures. (b) Dependence of $\mathcal{L}$ on $\chi$, showing a unique minimum.

### VI. DETERMINATION OF THE SURFACE TENSION

Here, we investigate the convergence of the estimated surface tension $\gamma$. We define a running average of each diagonal component $j$ of the pressure tensor using all measurements from $i = 0$ up to $i = n$:

$$\langle P_{jj} \rangle_n = \frac{1}{n+1} \sum_{i=0}^{n} P_{jj,i}. \tag{S2}$$

The corresponding running estimate of the surface tension is then given by

$$\gamma_n = \frac{L_z}{2} \left[ \langle P_{zz} \rangle_n - \frac{1}{2} \left( \langle P_{xx} \rangle_n + \langle P_{yy} \rangle_n \right) \right]. \tag{S3}$$

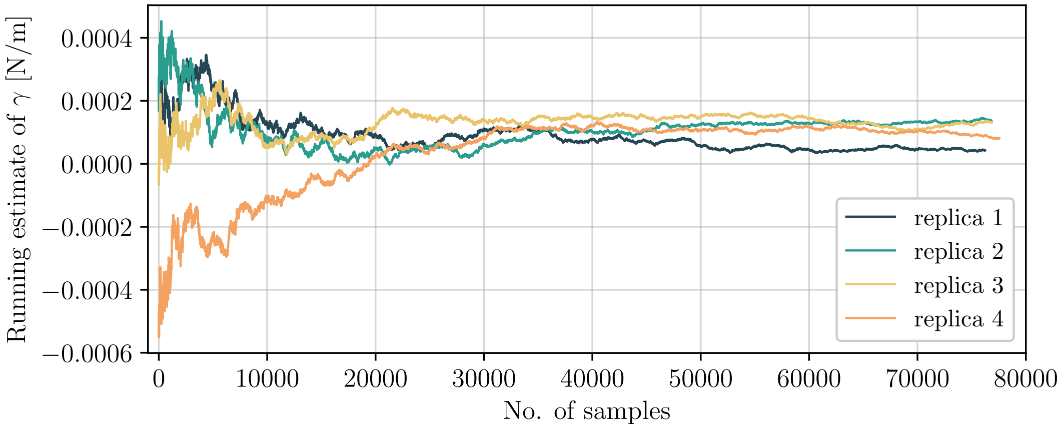


FIG. S5. Running average estimate of the surface tension for 4 independent replicas.

### VII. TIME-DEPENDENT STRUCTURE FACTORS DURING SPINODAL DECOMPOSITION

In Fig. S6, we provide an example of 15 time-dependent structure factors computed from MD trajectories and used to estimate $\omega(k)$. The dashed lines represent linear fits of $\log S(k,t)$ as a function of time. In the regression, we did not constrain the intercept to the value $\log S(k,0)$ predicted by the linearized Cahn–Hilliard theory. This is because the system may require a finite response time after the quench before entering the regime described by the linearized theory.

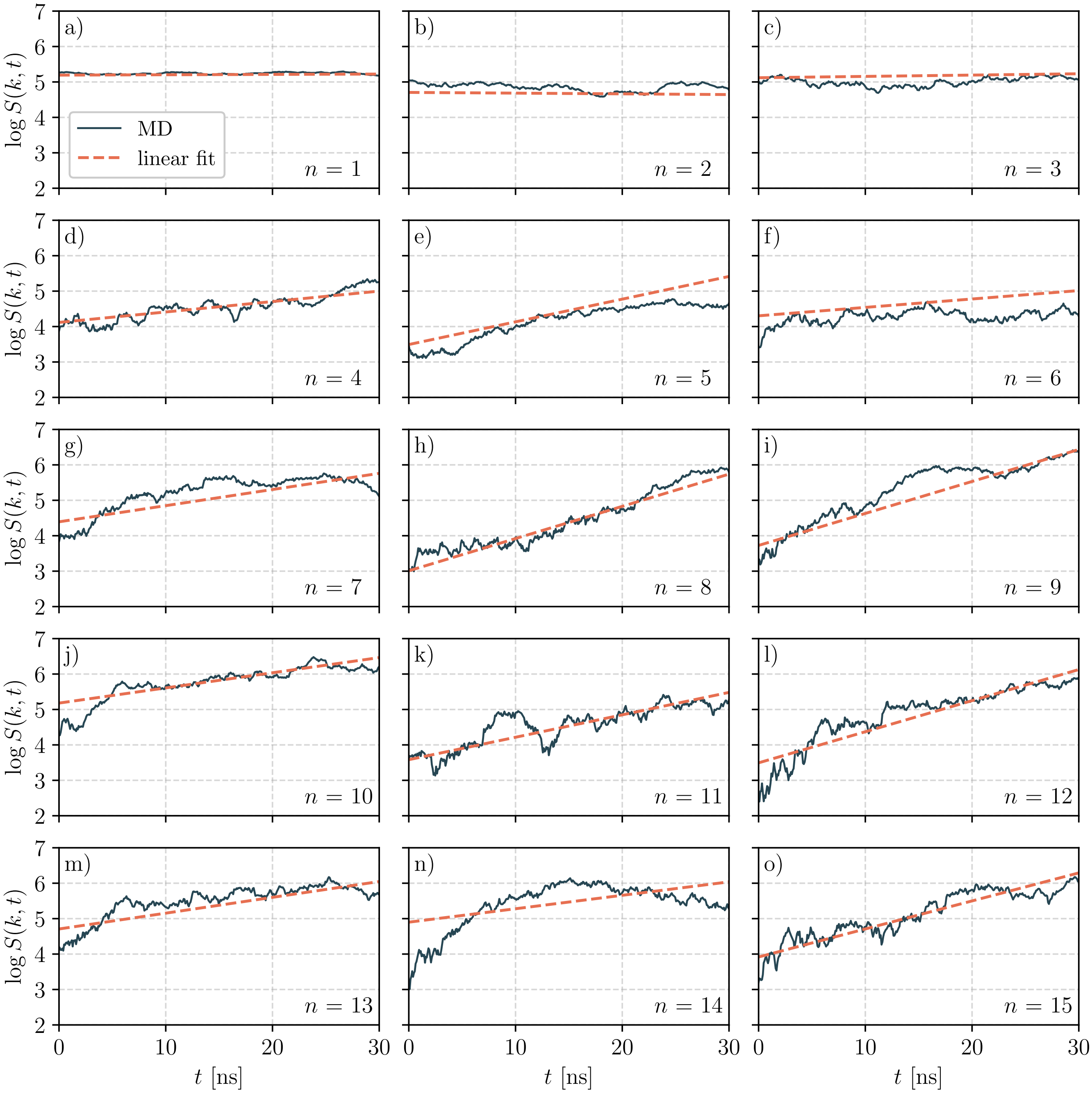


FIG. S6. (a)-(o) Time–dependent structure factors with the corresponding linear fits.

### VIII. DOMAIN FORMATION

Here, we show an example of domain formation as the system evolves from an initially uniform concentration and enters the coarsening stage.

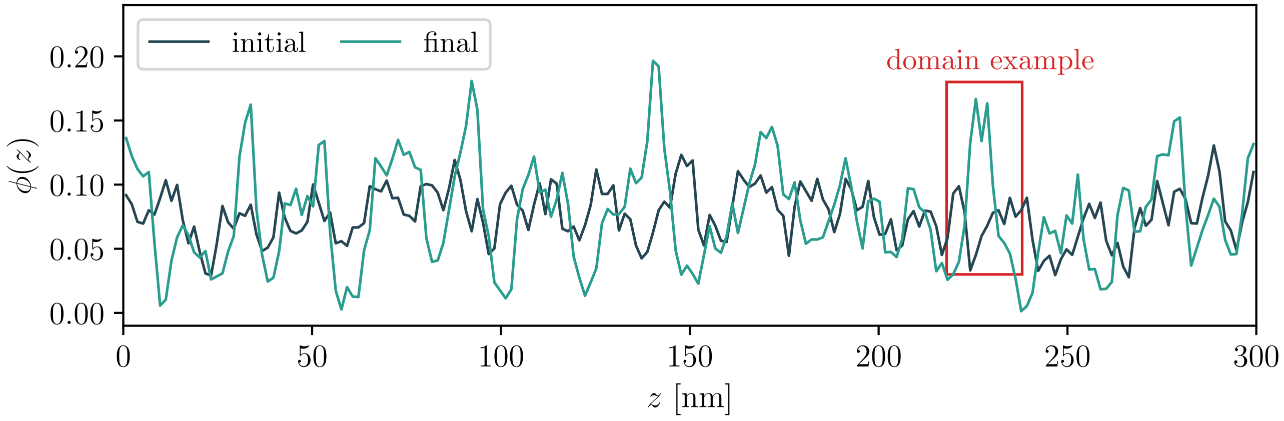


FIG. S7. Examples of domain formation within red rectangles as the system evolves for 30 ns from an initially uniform state.
